# Response of neuroglia to hypoxia-induced oxidative stress using enzymatically crosslinked hydrogels

**DOI:** 10.1101/799692

**Authors:** Samantha G. Zambuto, Julio F. Serrano, Avery C. Vilbert, Yi Lu, Brendan A.C. Harley, Sara Pedron

## Abstract

Three-dimensional cultures have exciting potential to mimic aspects of healthy and diseased brain tissue to examine the role of physiological conditions on neural biomarkers as well as disease onset and progression. Hypoxia is associated with oxidative stress, mitochondrial damage, and inflammation, key processes potentially involved in Alzheimer’s and multiple sclerosis. We describe the use of an enzymatically-crosslinkable gelatin hydrogel system within a microfluidic device to explore the effects of hypoxia-induced oxidative stress on neuroglia, astrocyte reactivity, and myelin production. This versatile platform offers new possibilities for drug discovery and modeling disease progression.

## 1. Introduction

There are multiple conditions that lead to abnormally low oxygen concentrations in tissues, such as hypoxemia in high altitude conditions, pulmonary disease, sleep disorders, blood vessel dysfunction that lead to ischemia, anemia and histotoxic hypoxia due to mitochondrial dysfunction^1^, and stroke. Many of these conditions may affect brain homeostasis and lead to neurological damage. Neurodegeneration is the main pathological characteristic of Alzheimer’s disease, Parkinson’s disease, stroke, traumatic brain injury and multiple sclerosis, among others. This devastating process commonly starts with the activation of microglia and the subsequent neuroinflammation may increase oxidative stress to neurotoxic levels. Studies have shown that oxidative stress caused by hypoxia can also induce inflammation followed by neuronal death and can be disrupted only by inhibiting inflammatory pathway components^2, 3^ However, moderate levels of hypoxia may improve resistance to later injuries caused by higher levels of hypoxia.^4^ Nevertheless, the neuroprotective – neurotoxic effects of hypoxia-induced oxidative stress on neural cells is not well understood.

Redox homeostasis is a determinant for the protein folding process^5^ and the production of reactive oxygen species (ROS), such as peroxides. These processes may activate integrated stress response (ISR), a common adaptative pathway that cells activate in response to different stress stimuli.^6^ However, the details regarding oxidative stress - ISR relationship in neuroglia are not well understood. Oxidative stress appears when the production of ROS surpasses the antioxidative capacity of the cell and can cause cell damage leading to apoptosis. Hypoxia increases the presence of ROS independently of hypoxia-inducible factor (HIF).^7^ Hypoxia is thought to facilitate pathogenesis of certain neurodegenerative disorders. For example, the incidence of Alzheimer’s disease increases following conditions that cause significant cerebral ischemia, such as stroke, obstructive sleep apnea syndrome and chronic obstructive pulmonary disease.^8, 9^ Amyloid precursor protein (APP) levels have been shown to be elevated in human brains after ischemic conditions which is thought to result in increased deposition of amyloid β, a hallmark of this disease.^8^ There is also evidence that hypoxia/inflammation cycles are associated with multiple sclerosis, a neurodegenerative disease characterized by a loss of myelin from inflammation. Clinical trials have demonstrated that individuals with multiple sclerosis suffer from reduced cerebral blood flow which likely causes hypoxic stress in the brain.^10^ This hypoxic environment is thought to form a positive-feedback loop with inflammation which further damages blood vessels and results in a chronic hypoxic/inflammatory condition that continues to damage cells in the brain.^10^ Nevertheless, the mechanisms regarding how hypoxia contributes to pathogenesis, progression, and onset of neurodegenerative disorders remain poorly understood. The analysis of neural toxicity and damage induced by metabolic dysfunction would provide a better understanding of tissue damage and identify more objective predictive biomarkers that will aid in diagnosis and treatment of disease.

Experimental access to key cell types in the nervous system to understand their roles in brain function and response to injury provides a challenge to understanding human neural function.^11^ Rodent models have provided a better understanding of neurodevelopmental disorders and development of more efficient treatments; however, the translation of *in vivo* results into human clinical trials has often been unsatisfactory.^12, 13^ Some *in vitro* strategies, such as organoids, have helped in reducing the gap between model and clinical practice but the complex cellular composition of organoids has led to low reproducibility and the lack of vascularization in organoids limits oxygen availability in some parts of the tissue.^12, 13^ Advanced biomaterial platforms enable researchers to mimic the three-dimensional (3D) neural architecture of the brain, heterogeneous population of cells (*e.g.*, neurons, astrocytes, oligodendrocytes, microglia), and extracellular matrix environment (*e.g.*, stiffness, matrix composition) in reproducible, high throughput systems. Such models provide researchers with opportunities to investigate underlying pathological mechanisms of neurodegenerative disorders in complex, 3D platforms that can be tuned to replicate the diseased microenvironment.^14, 15^ In this study, we employ enzymatically-crosslinked hydrogels that are able to consume oxygen during the polymerization reaction, creating a controllable temporary hypoxic environment that leads to the production of ROS. Furthermore, the use of an analog reaction that uses hydrogen peroxide as substrate instead of oxygen affords a different source of oxidative stress, in addition to the appropriate modification of mechanical properties. This miniaturized system maintains culture of neural stem cells and the regulation of their differentiation into relevant cell types. These hydrogels provide a controlled environment and bioactive milieu adapted to study hypoxia-induced oxidative stress. The results of our studies presented herein may provide deeper insights into the role of hypoxic environment on the onset and progression of the disease, assisting in the development of novel therapeutics to repair damaged areas.

## 2. Materials and Methods

### Materials

Gelatin (type A from porcine skin, less than 300 bloom), 3-methoxy-4-hydroxycinnamic acid (ferulic acid, FA), 3-(ethyliminomethyleneamino)-*N*,*N*-dimethylpropan-1-amine hydrochloride (EDC), *N*-hydroxysuccinimide (NHS), dimethyl sulfoxide (DMSO), and deuterium oxide (D_2_O) were purchased from the Sigma-Aldrich Company (Saint Louis, MO) and used without further purification. Purified laccase from *Streptomyces coelicolor* (aqueous solution of 192 U/mL) was obtained as a gift from Prof. Yi Lu (University of Illinois). Pierce™ Horseradish peroxidase (HRP) was purchased from Thermofisher scientific. Phosphate-Buffered Saline (PBS), Dulbecco’s Modified Eagle Medium (DMEM), and fetal bovine serum (FBS) were all purchased from Gibco, Invitrogen (Life Technologies, CA). Hoechst and CellROX™ Green Reagent were purchased from Thermo Fisher Scientific. Dialysis membrane (molecular cutoff 3.5 kDa) was purchased from Spectrum Laboratories (Rancho Dominguez, CA). 3D Cell Culture Chips were purchased from AIM Biotech (Singapore).

### Instrumentation

Nuclear magnetic resonance (NMR) spectra were recorded on a Varian Unity 500 at ambient temperature. Chemical shifts (δ) are reported in parts per million (ppm). ^1^H NMR chemical shifts were referenced to the residual solvent peak at 4.80 ppm in D_2_O. Compressive moduli analyses were performed on an Instron 5943 Single Column Tabletop Testing System (Eden Prairie, MN). Hydrogels were analyzed using confocal microscopy (Zeiss LSM710).

### 2.1. Hydrogel fabrication and characterization

Gelatin-ferulic acid (GelFA)^16^ and gelatin-hydroxyphenylpropionic acid (GelHPA)^17^ were synthesized according to a procedures described in literature. Resulting products were characterized by NMR and UV absorbance at 320 nm to account for conjugated ferulic acid molecules. Hydrogel precursor solution were prepared in DPBS. Laccase final concentrations varied from 25 U/ml to 10 U/ml and were used to fabricate GelFA hydrogels with concentrations between 5 and 3wt%. Horseradish peroxidase (HRP) concentrations were 0.1 and 0.25 U/ml and reacted in conjunction with 1 - 2 mM hydrogen peroxide solution respectively. HRP catalyzed the fabrication of GelFA (3wt%) and GelHPA (2wt%) hydrogels. Gelatin conjugated with ferulic acid and hydroxyphenylpropionic acid groups acts as a hydrogen-donating substrate (AH). Laccase reduces oxygen while concurrently oxidizing ferulic acid (4AH) and HRP catalyzes the reduction of hydrogen peroxide by oxidizing two equivalents of AH (2AH). The hydrogen subtraction leads to formation of C and O radicals in the phenoxy group that subsequently form the crosslinking bonds (C-C and C-O) leading to hydrogel structure.

Compressive Young’s moduli were obtained from the linear region of the stress–strain curve (15% strain) as previously described.^18^ Hydrogel samples of the desired wt% were fabricated using Teflon molds (0.2 mm thick, 10 mm diameter) and allowed to swell in PBS for 16 h at 37°C. Samples were compressed at the rate of 0.1 mm/min. Moduli were calculated using a custom MATLAB code, which calculates modulus from the linear regime at a load of 0.03 N and offset of 2.5% to ensure contact with the hydrogel surface.

**Table 1.**
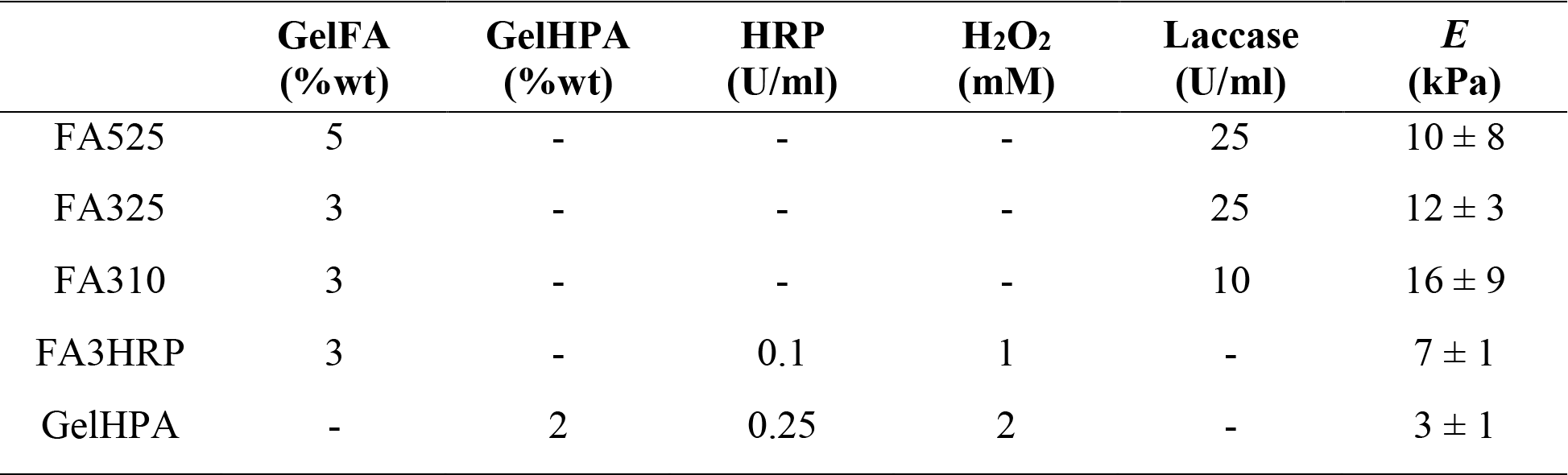
Hydrogel composition, polymerization conditions and Young’s moduli.

Oxygen partial pressure was quantified using Oxoplate OP96C (Precision Sensing GmbH, Regensburg, Germany). Hydrogels were prepared within the bottom of 96-well plate in the presence of an oxygen sensor and within a regular 96-well plate (no sensor), used as control. Fluorescence was recorded in a plate reader (Biotek Synergy HTX) at ex540nm/em650nm and ex540nm/em650nm following the manufacturer’s protocol 16 h after polymerization occurred. Additionally, fluorescence background from the regular plate was subtracted from values obtained from the sensor prior to calculating % pO_2_.

### 2.2. Cell Culture and encapsulation

Normal human astrocytes (NHA, Lonza; CC-2565), rat fetal neural stem cells (NSC, ThermoFisher; N7744100) and mouse oligodendrocytes (OD, Celprogen; 11004-02) were used for experiments. Cells were maintained as per the manufacturer’s instructions in growth medium supplemented with 1% penicillin/streptomycin and were used no more than 4 passages of NHA, 5 passages of OD and 6 passages of NSC after purchase. Information regarding cell media formulations can be found in Supplemental Table S1. Cells were cultured in 5% CO_2_ incubators at 37 °C. Routine mycoplasma testing was performed on the cells every 6 months.

#### Preparation of cell laden hydrogels within microfluidic chips

GelFA was dissolved in DPBS at 37 °C. A solution of laccase was prepared with a final activity concentration of 100 U/ml. Cell pellets were resuspended using the GelFA solution (2×10^6^ cells/ml NHA, 20×10^6^ cells/ml NSCs, 4×10^6^ cells/ml ODs final concentration) before addition of laccase solution (final concentration 3-5 wt% GelFA and 10-25 U/ml laccase) and injection into the cell culture chips. After 7 min of injection, medium was added to the media channels of the cell culture chips. The chips were incubated at 37°C, 5% CO_2_ and controlled humidity. For GelFA hydrogels using HRP and H_2_O_2_, GelFA was dissolved in DPBS, mixed with a solution of HRP, and used to resuspend cell pellets (20×10^6^ cells/ml NSCs, 4×10^6^ cells/ml ODs final concentration). Then, a solution of H_2_O_2_ was added and quickly mixed (final concentration 3 wt% GelFA, 0.1 U/ml HRP, and 1 mM H_2_O_2_) and injected into the middle channel of the cell culture chips. The appropriate medium was added after 5-7 min to allow for polymerization, and the same steps for incubation as described aboved were followed. Chips were maintained in cell growth medium and medium was replaced daily.

GelHPA was dissolved in DPBS buffer at 37 °C and mixed with HRP solutions (final concentration 2 wt% GelHPA and 0.25 U/ml). Cell pellets were resuspended (4×10^6^ cells/ml ODs, 2×10^6^ cells/ml NHA final concentration) using this solution to give *ca*. 100 μl of solution, and H_2_O_2_ (2 mM) was added immediately before injection into the cell culture chips. Each aliquot was used to seed one chip to ensure a homogeneous gel formation across all channels. Seven minutes following injection, medium was added to the media channels of the cell culture chips. Chips were maintained in cell growth medium and medium was replaced daily.

Cell differentiation was achieved following NSC’s manufacturer’s (ThermoFisher Scientific) protocol. Proliferation and differentiation media compositions are included in the Supplementary Information.

HRP polymerized samples were cultured under 1% O_2_ atmosphere in a computer-controlled hypoxia chamber (C-Chamber Hypoxia Chamber, Biospherix™, Parish, NY) to induce a hypoxic environment.

### 2.3. Qualitative analysis of reactive oxygen species production

Oxidative stress was determined using CellROX Green Reagent, a fluorescent probe for detecting cellular oxidative stress. Samples were incubated for 1 h in 5 μM CellROX Green diluted in sterile cell growth medium. After 1 h, the CellROX solution was discarded and three PBS wash steps were performed. Samples were fixed in Image-iT Fixative Solution for 15 min followed by three PBS washes, stained with Hoechst (1:2000) for 30 min, washed once with PBS, and stored in PBS until imaged, using DMi8 Yokogawa W1 spinning disc confocal microscope outfitted with a Hamamatsu EM-CCD digital camera (Leica Microsystems) or confocal microscope Zeiss LSM 710. Representative images are shown for n=3 hydrogels.

### 2.4. Immunofluorescent staining

Information regarding antibody dilutions can be found in the Supplementary Information Table S2. Hydrogels were fixed in Image-iT Fixative Solution for 15 min followed by three PBS washes. Cells were permeabilized in a 0.1% Triton-X-100 solution in PBS for 15 min at room temperature followed by three five-minute PBS washes. Samples were blocked for 2 h in a 2% bovine serum albumin (BSA) solution in PBS and subsequently incubated in the primary antibody solution diluted in a 2% BSA solution overnight at 4°C. Following five five-minute PBS washes, samples were incubated in the secondary antibody solution in a 2% BSA solution overnight at 4 °C. After five five-minute PBS washes, samples were incubated for 30 min at room temperature in a 1:2000 dilution of Hoechst in PBS. Samples were washed one time in PBS and stored in PBS at 4 °C until imaged using confocal microscope Zeiss LSM710. Representative images are shown of n=3 hydrogels. GFAP fluorescence was quantified using ImageJ. The corrected total cell fluorescence was calculated measuring the area integrated fluorescence intensity in 10 points per image relative to background readings for same area.

### 2.5. Statistics

OriginPro 2018b (Origin Lab) or RStudio were used for all statistical analysis. The Shapiro-Wilkes test was used to determine normality and Levene’s test was used to determine equality of variance. Mechanical testing data was analyzed using a Kruskal-Wallis one-way analysis of variance (ANOVA) with Dunn’s post-hoc analysis for n=6-9 hydrogels per condition. Mechanical testing data for FA525 and GelHPA conditions were analyzed via a two-way t-test with Welch’s correction. Oxygen concentration was calculated from n=4 samples using Welch’s ANOVA followed by Games-Howell post-hoc pairwise mean comparison (p<0.05). Corrected total cell fluorescence (GFAP) is shown for n=3 hydrogels (normoxia vs hypoxia) and significance is calculated using one-way analysis of variance ANOVA followed by Tukey’s HSD post-hoc test (p<0.05). Significance was set at p < 0.05. Results are listed as mean ± standard deviation unless otherwise noted.

## 3. Results and discussion

### 3.1. Stiffness and oxygen content regulate stem cell phenotype

The nature of gelatin-based enzymatically crosslinked hydrogels allows us to manipulate physical properties of the stem cell microenvironment along with the oxygen concentration. Both properties are important in controlling stem cell behavior within three-dimensional materials, such as proliferation, cell spreading, and subsequent differentiation into neural lineages. Cells were cultured within microfluidic chips (**Figure 1a**) that include a dedicated central channel for the formation of the 3D hydrogel, bordered by posts to retain the hydrogel zone between two flanking media channels. We first examined the biophysical properties of a series of gelatin hydrogel networks that relied on either more conventional HRP mediated crosslinking (GelHPA macromer) that has previously been shown to generate radicals, or a laccase-mediated reaction (GelFA macromer) that consumes oxygen. Gelatin was used as a constant macromer backbone (at 3mg/ml) as it provides cell adhesion and degradation sites. We observed altering laccase concentration (10 vs. 25 U/ml laccase) primarily affects oxygen availability but does not significantly affect resultant modulus. Laccase polymerized hydrogels (FA325, FA310) displayed Young’s moduli (*E*) between 12±3 and 16±9 kPa, while HRP polymerized gelatin hydrogels (FA3HRP) displayed a significantly softer (7±1 kPa) hydrogel environment. Statistical analysis (Kruskal-Wallis ANOVA) revealed significant differences amongst groups (Chi square = 11.8, *p* = 0.0027, df = 2) with HRP crosslinked hydrogels significantly softer than the laccase gelation process as indicated by Tukey HSD post-hoc analysis (HRP compared to 25U and 10U laccase; *p* = 0.014 and *p* = 0.0096 respectively).

**Figure 1.**
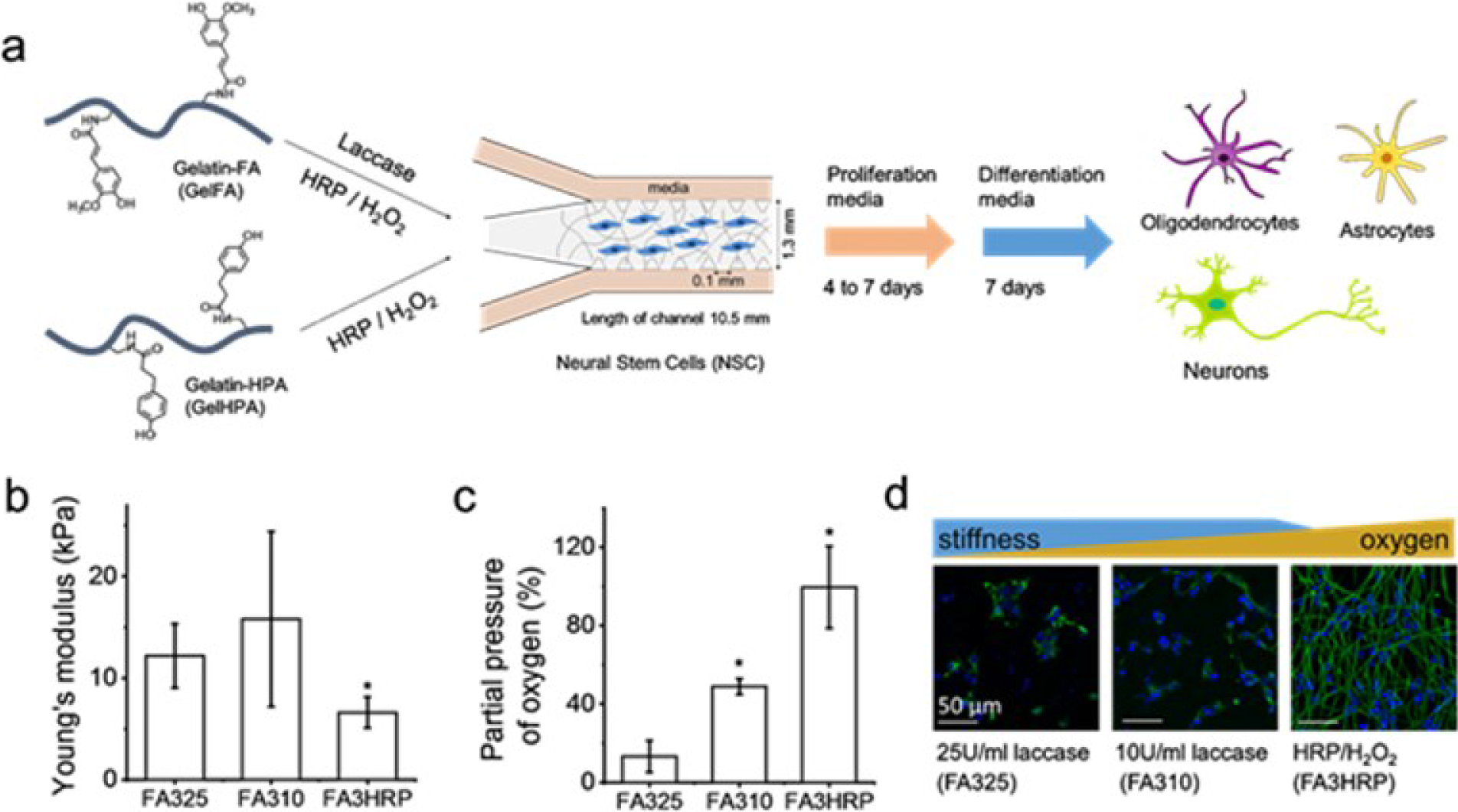
Neural stem cells maintain their phenotype in enzymatically-polymerized gelatin hydrogels. (a) Hydrogels based on GelFA and GelHPA are formed in the presence of laccase or HRP/H_2_O_2_ within AIM Biotech microfluidic chips. Differentiation was induced after 4 or 7 days in maintenance media to allow for matrix remodeling. (b) Mechanical properties (*p<0.05 compared to FA3 samples). (c) %pO_2_ varied depending on enzyme concentration, %wt and enzyme type. Mean ± SD, *p<0.05 compared to FA325. (d) Rat neural stem cells are stained for Nestin (green) and Hoechst (blue) within 3wt% GelFA hydrogels.

The method of hydrogel gelation significantly impacted the resultant partial oxygen pressure within the hydrogel (**Figure 1c**). Notably, significantly reduced oxygen availability with increasing amount of laccase used to catalyze hydrogel gelation (oxygen partial pressure: 13±8% for 25U/ml, 49±4% for 10U/ml, *p*=0.01) was significantly lower than that for conventional HRP/H_2_O_2_ mediated crosslinking (100±21 %pO2 HRP/H_2_O_2_). There was a significant effect on oxygen concentration based on the gel formulations at the p<0.001 level for the three conditions (p=0.0009). Games-Howell post-hoc analysis indicated that all three groups (FA325, FA310, FA3HRP) are statistically significantly different from each other (*p*<0.05). Notably, increasing the gelatin concentration to 5wt% did not further increase the elastic modulus of laccase crosslinked hydrogel (*E*=10±8 kPa, *p*=0.97 one-way ANOVA). NSCs cultured in maintenance medium within these hydrogels did not exhibit morphological shifts associated with spontaneous differentiation. Cells show expression of Nestin, a stem cell marker, in immunostaining experiments for all conditions. However, softer hydrogel environments (FA3HRP) supported increased NSC spreading of cells. The reduced oxygen availability in FA hydrogel variants may explain increased cell aggregation in the FA325 hydrogel variant that displayed the lowest oxygen availability, a phenomenon previously described for endothelial cells.^19^

### 3.2. Enzymatically polymerized gels support NSC differentiation

Neural stem cells hold a great potential to model healthy and diseased brain tissue. Multiple platforms have been designed to maintain and induce differentiation based on mechanical properties,^***20***^ degradability^***21***^ and ligand presentation,^***22***^ among others. Previous studies have shown that softer matrices, between 0.5 and 1.5 kPa, support NSC differentiation into neurons and astrocytes.^***23***^ As a result, here we examined NSCs differentiation patterning into multiple neural cell types, notably neurons, astrocytes, and oligodendrocytes as a result of exposure of NSCs to neuronal or astrocytic differentiation media. We observed GelFA hydrogels are able to maintain stem cells in different degrees depending on stiffness and oxygen concentration. As matrix remodeling has been reported as a requirement for NSC to effectively differentiate,^24^ we also examined whether differentiation efficiency shifted after 4 or 7 days in proliferation media prior to differentiation induction. Immunostaining images of astrocyte marker glial fibrillary acidic protein (GFAP) and neuronal marker β–III-tubulin (**Figure 2a**) show that softer materials support effective neuronal differentiation, while astrocyte phenotype but no neural phenotype is observed in stiffer matrices that exhibited early levels of laccase-mediated hypoxia during gelation (FA325). Astrocyte cell spreading, however, was more pronounced in laccase polymerized gels with higher %pO2. Culture time prior to differentiation (4 days versus 7 days) affects astrocyte differentiation efficiency in softer HRP matrices, probably associated to an increased matrix remodeling.^***24***^ Together, these hydrogels provide a suitable platform for differentiation of both astrocytes and neurons, resulting in interconnected networks of neurons and neuroglia. The use of co-cultures may allow the study of the function of local microenvironment in neural damage and the role of glial cells in neuroprotection. We are interested in stages of brain disease where oxygen availability is hindered, for instance during inflammation. Those stages are important in demyelination processes. Therefore, we also evaluated mature oligodendrocyte differentiation (O1 marker) and secretion of myelin basic protein (MBP), a protein that maintains the structure of myelin and that is expressed exclusively in myelinating glia (**Figure 2b**). We only observed effective OD differentiation in softer matrices. Furthermore, later differentiation induction (7 days) provided a more robust differentiation of oligodendrocytes, and the presence of this platform in a hypoxic environment (1% O_2_ hypoxic chamber) did not alter MBP expression (**Figure 2c**).

**Figure 2.**
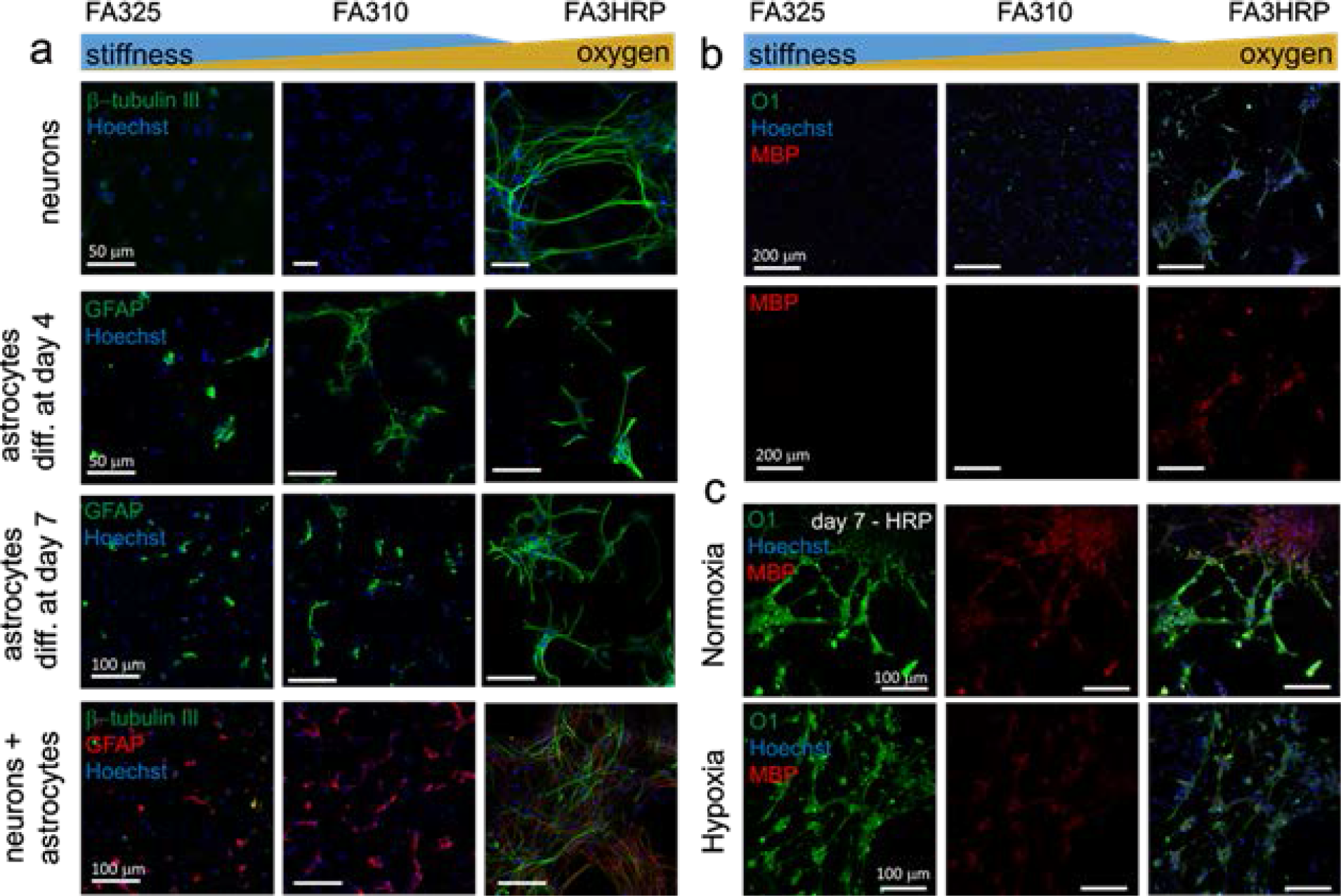
Neural Stem Cell differentiated into astrocytes, neurons and oligodendrocytes. (a) Stiffness and oxygen concentration influence the extent of cell differentiation into astrocytes (GFAP) and neurons (β-tubulin III). Differentiation into astrocytes was induced at day 4 and day 7. (b) Stiffness and oxygen concentration influences NSC differentiation into mature oligodendrocytes (O1) and expression of myelin (MBP). Hypoxia (1% O_2_, for 48h) environment was applied to HRP-polymerized hydrogels differentiated into oligodendrocytes. 3wt% GelFA is used for all hydrogels, 25 and 10 U/ml of laccase and 0.1U/ml HRP – 1mM H_2_O_2_.

### 3.3. Astrocytes become reactive under oxidative stress

Hypoxia induces the production of reactive oxygen species (ROS), that are released by activated microglia through respiratory burst that produces radicals and hydrogen peroxide into the extracellular matrix.^25^ The precise role of astrocytes in degeneration is still unclear, they may play a role in demyelination processes by affecting inflammation, leading to cell damage and glial scarring.^26^ Astrocytes are believed to have a high tolerance to oxidative stress during pathological situations.^27^ Moreover, astrocytes exposed to oxidative stress become preconditioned against repetitive exposure.^27^ It has been reported that acute hypoxia induces oxidative stress by increasing the production of ROS in the brain due to the alteration of the mitochondrial oxidative phosphorylation. Hydrogen peroxide is a widely-known member of the ROS family that can modulate signaling pathways, inducing apoptosis but can also stimulate proliferation at low concentrations. Using the suite of laccase and HRP mediated crosslinked hydrogels, we were able to examine the role of oxidative stress on astrocyte activity where oxidative stress could be induced by: ***(1)*** exposure of astrocyte-hydrogel cultures to a low oxygen atmosphere (1% O_2_) for 24 h; ***(2)*** during HRP-mediated crosslinking of GelHPA hydrogels due to exposure to H_2_O_2_; or ***(3)*** exposure to a hypoxic microenvironment transiently created by laccase (an oxygen-consuming enzyme) mediated GelFA hydrogel crosslinking.

We examined ROS formation within the hydrogel microenvironment using CellROX™ (**Figure 3**), a dye that becomes fluorescent in the presence of ROS and binding to DNA. Astrocytes in GelHPA hydrogels polymerized with hydrogen peroxide show significantly higher ROS levels under normoxic culture than hypoxic culture (**Figure 3a**). Interestingly, ROS levels in GelHPA hydrogels are maintained for at least 48 h, and at levels higher than GelHPA cultures exposed to 24 hours of hypoxia followed by 24 hours of normoxia (**Figure 3b**), likely due to increased formation of oxygen derived radicals during hydrogen peroxide mediate hydrogel crosslinking. On the other hand, astrocytes in laccase crosslinked GelFA hydrogels (**Figure 3c**) display transient production of ROS 1h after polymerization that disappears at 24h. Cell morphology differences in GelFA/laccase versus GelHPA/HRP platforms may be due to differences in mechanical properties (10±8 versus 3±1 kPa respectively,; *p*=0.030 Student’s t-test with Welch’s correction). While we have demonstrated astrocytes become reactive in response to pathological situations, little is known about their effect in the progression of different neurodegenerative diseases. Pathological studies in AD diseased tissue samples have demonstrated the presence of reactive astrocytes, characterized by an increased expression of GFAP.^28^ Astrocytes cultured in HRP/H_2_O_2_ crosslinked GelHPA hydrogels show significant (*p*=5E-4 between normoxia and hypoxia samples) increases in GFAP expression after 24h in hypoxia (**Figure 3d**), indicating that reactivity of astrocytes is independent of the production of oxygen derived radicals during hydrogel formation.

**Figure 3.**
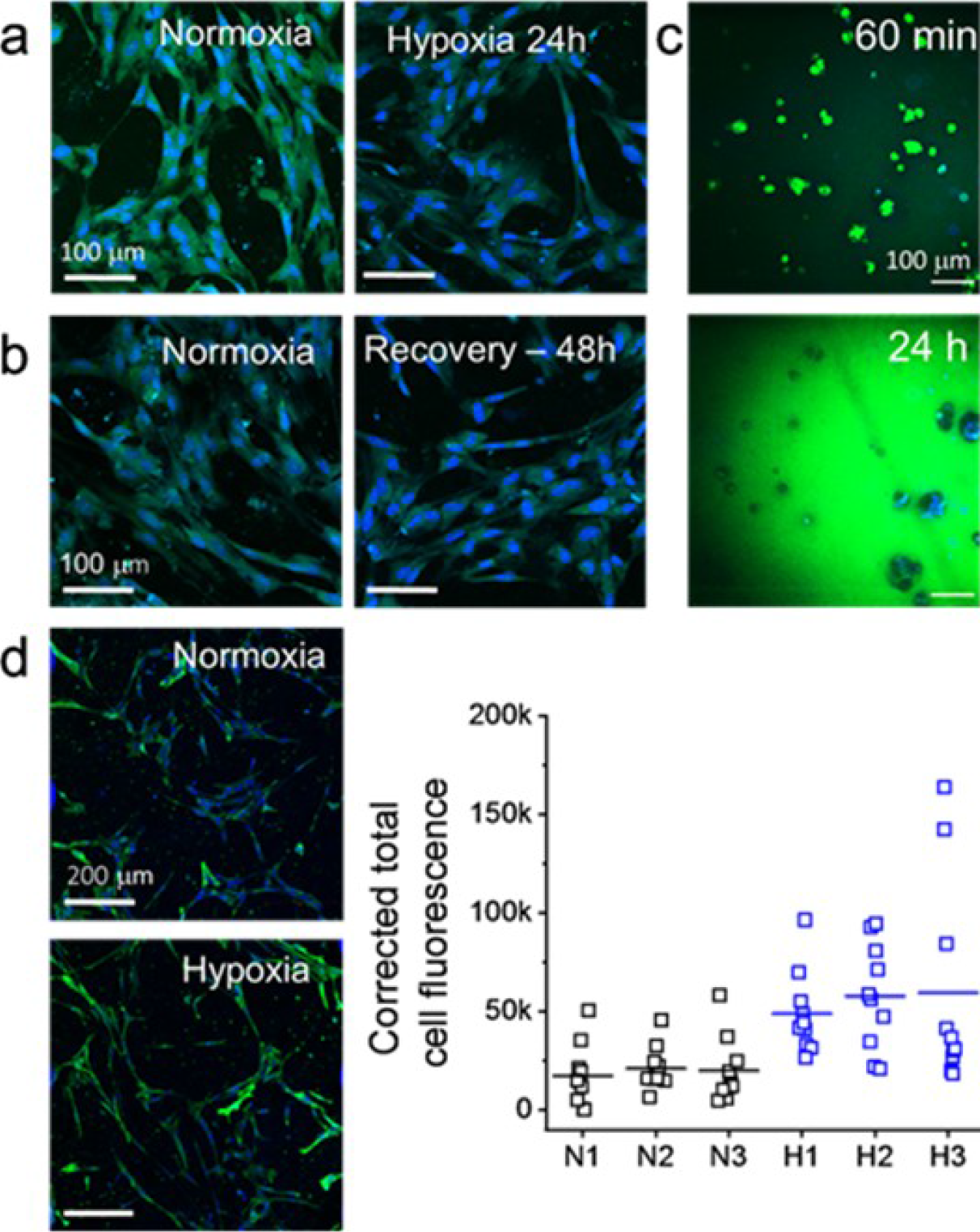
Astrocytes become reactive under hypoxic conditions. (a) Reactive oxygen species (ROS) analyzed by CellROX (green) are detected in astrocytes within 2wt% GelHPA hydrogels in normoxia, after 24h in hypoxia (1%O_2_) and (b) following 24h of recovery in normoxia (as compared to continuous normoxic conditions). 2wt% GelHPA hydrogels are prepared with HRP/H_2_O_2_. (b) CellROX analysis of ROS in 3wt% GelFA hydrogels / laccase 25U/ml at different time points. (d) GFAP (green) expression of astrocytes increases in hypoxic (1%O_2_) environment. Results of corrected total fluorescence between N (normoxia) and H (hypoxia) samples are significantly different p<0.05.

### 3.4. Hypoxia does not affect myelin production in mature oligodendrocytes

Vascular disease and metabolic dysfunction caused by toxins released by activated microglia may be responsible of demyelinated lesions.^29^ Oligodendrocytes (ODs) are the myelin-forming cells, they are present in the brain in all forms from pre-progenitors to mature oligodendrocytes. Therefore, they are crucial in the maintenance and remyelination after damage of the myelin sheath that forms around the nerves. The ability to manipulate ODs in diseased microenvironment may aid the development of more effective remyelination strategies.^30^ Strategies addressed to promote myelination show recovery of neuronal function caused by low oxygen in brain tissue.^31^ The understanding of neurodegeneration in inflammatory demyelinating diseases, such as multiple sclerosis, requires more appropriate models that reflect OD response and myelin secretion during disease progression. We have shown that oxygen deprivation for 48 h does not significantly change MBP expression of differentiated oligodendrocytes (**Figure 2c**). Additionally, we have cultured embryonic oligodendrocytes within 3wt% GelFA hydrogels. We showed that ODs in less crosslinked hydrogels (**Figure 4b**) show larger spheroids than in stiffer ones (**Figure 4a**). Previous reports indicated that oligodendrocyte progenitor cells have a preference for proliferation in softer materials in detriment of differentiation into mature oligodendrocytes.^32^ Likewise, it has also been described that physical properties and mechanisms associated with hypoxia upregulate proteases, through production of ROS, leading to cell aggregation.^19^ Here, we also showed that secretion of MBP is not dependent on low oxygen concentration (hypoxia chamber for HRP gels) or laccase induced hypoxia. As a result, *ex vivo* biomaterial models may provide the opportunity to examine the processes that lead to OD damage such as oxidative stress, where chronic hypoxia may induce myelin loss and neurodegeneration.

**Figure 4.**
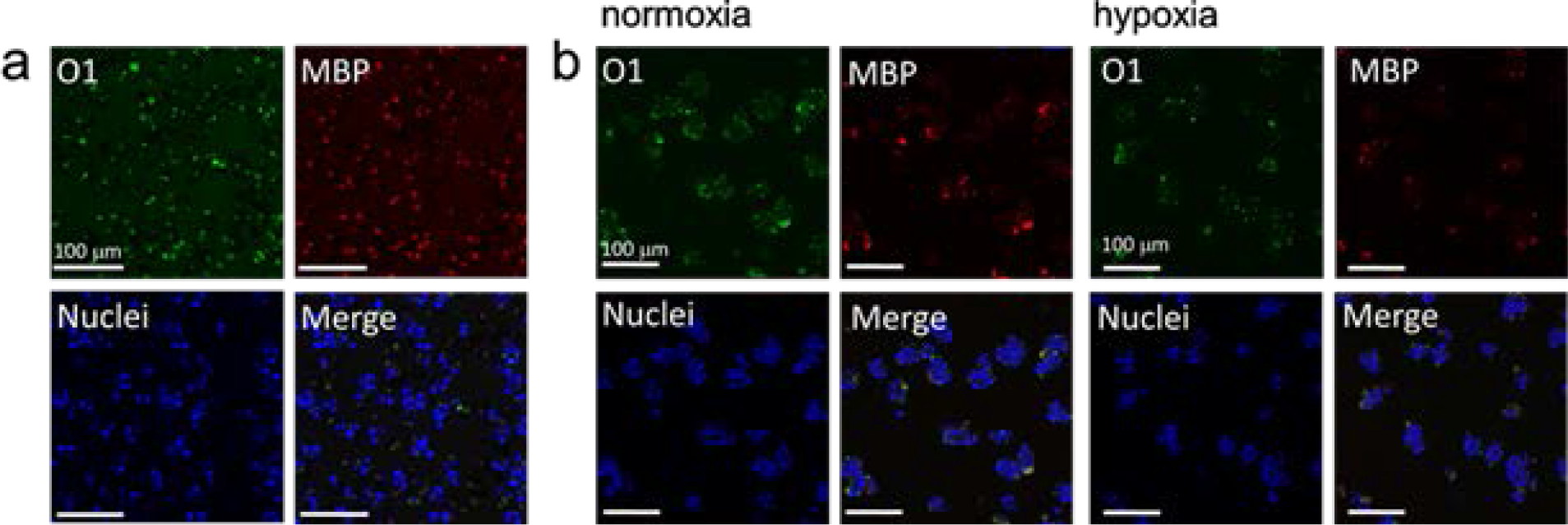
Effect of hypoxia in mature oligodendrocytes and myelin expression. (a) ODs cultured within 3wt% GelFA / laccase 25U/ml (FA325) and (b) 3wt% GelFA / HRP/ H_2_O_2_ hydrogels (FA3HRP) stained for O1 (green, mature OD marker) and MBP (red, myelin basic protein). HRP-polymerized hydrogels were exposed to hypoxia (1%O_2_) for 24h.

## 4. Conclusions

The advent of advanced biomaterial-based cell culture platforms has allowed researchers to create models of the brain with increasing complexity. Such platforms are urgently required to mitigate poor translation of *in vitro* and animal models to human physiology and disease. We demonstrated hydrogel platforms that are able to maintain and differentiate neural cells into relevant linages and evaluate neuroglia reaction to disparate levels of hypoxia. Notably, the use of laccase-mediate crosslinking provides an avenue to induce transient levels of hypoxia within the hydrogel sufficient for ex vivo examination of the role of hypoxic stress on neural cell populations. These induced environments may be of significant interest to evolving studies of neurodegenerative diseases.

## Acknowledgements

The authors would like to acknowledge Dr. Austin Cyphersmith (IGB, University of Illinois) for assistance with fluorescence imaging, Mai Ngo for assistance with microfluidic chips, and Zona Hrnjak and Aidan Gilchrist for the compression testing methodology and the custom MATLAB code for modulus analysis. The authors gratefully acknowledge support from the Scott H. Fisher IGB Graduate Student Research Fund. Research reported in this publication was supported by the National Cancer Institute of the National Institutes of Health under Award Number R01 CA197488 as well as National Institute of Diabetes and Digestive and Kidney Diseases of the National Institutes of Health under Award Number R01 DK099528. The content is solely the responsibility of the authors and does not necessarily represent the official views of the NIH. The authors are also grateful for additional funding provided by the Department of Chemical & Biomolecular Engineering and the Carl R. Woese Institute for Genomic Biology at the University of Illinois at Urbana-Champaign.

