## Supplementary Information for "Response of neuroglia to hypoxia-induced oxidative stress using enzymatically crosslinked hydrogels"

^3^ Dept. Chemical and Biomolecular Engineering, University of Illinois at Urbana-Champaign, Urbana, IL

^4^ Pacific Northwest National Laboratory, Richland, WA

^5^ Dept. of Chemistry, University of Illinois at Urbana-Champaign, Urbana, IL

*These authors contributed equally to this work

**Co-corresponding authors

**Corresponding Authors:**

S. Pedron

Dept. of Chemical and Biomolecular Engineering

Carl R. Woese Institute for Genomic Biology

University of Illinois at Urbana-Champaign

1260 W Gregory Dr.

Urbana, IL 61801

B.A.C. Harley

Dept. of Chemical and Biomolecular Engineering

Carl R. Woese Institute for Genomic Biology

University of Illinois at Urbana-Champaign

110 Roger Adams Laboratory

600 S. Mathews Ave.

Urbana, IL 61801

**Table S1.** Hydrogel composition, polymerization conditions and Young’s moduli.

|  | GelFA  (%wt) | GelHPA  (%wt) | HRP  (U/ml) | H_2_O_2_  (mM) | Laccase  (U/ml) | *E*  (kPa) |
| --- | --- | --- | --- | --- | --- | --- |
| FA525 | 5 | - | - | - | 25 | 10 ± 8 |
| FA325 | 3 | - | - | - | 25 | 12 ± 3 |
| FA310 | 3 | - | - | - | 10 | 16 ± 9 |
| FA3HRP | 3 | - | 0.1 | 1 | - | 7 ± 1 |
| GelHPA | - | 2 | 0.25 | 2 | - | 3 ± 1 |

**Table S2.** Cell media composition.

| **Medium Type** | **Formulation** | **Concentration** |
| --- | --- | --- |
| Rat Fetal Neural Stem Cell Growth Medium | KnockOut DMEM/F-12  GlutaMAX-I Supplement  bFGF (prepared as 100 μg/mL stock)  EGF (prepared as 100 μg/mL stock)  StemPro Neural Supplement  Penicillin/Streptomycin | 1X  2 mM  20 ng/mL  20 ng/mL  2%  1% |
| Rat Fetal Neural Stem Cell Neural Differentiation Medium | Neurobasal Medium  B-27 Supplement (50X), serum free  GlutaMAX-I Supplement  Penicillin/Streptomycin | 1X  2%  2 mM  1% |
| Rat Fetal Neural Stem Cell Astrocyte Differentiation Medium | DMEM  N-2 Supplement  GlutaMAX-I Supplement  FBS  Penicillin/Streptomycin | 1X  1%  2 mM  1%  1% |
| Rat Fetal Neural Stem Cell Oligodendrocyte Differentiation Medium | Neurobasal Medium  B-27 Supplement (50X), serum free  GlutaMAX-I Supplement  T3  Penicillin/Streptomycin | 1X  2%  2 mM  30 ng/mL  1% |
| Normal Human Astrocyte Medium | ABM Basal Medium (Lonza CC-3187)  AGM Single Quots Supplement Pack (CC-4123) | 1X  1X |
| Mouse Embryonic Oligodendrocyte Medium | Mouse Oligodendrocytes Cell Culture Complete Media with Serum.  Cat# M11004-02 – Celprogen Inc. | 1X |

**Table S3.** Complete list of antibodies used for immunofluorescent staining.

| **Antibody** | **Vendor** | **Part Number** | **Host Species** | **Dilution** |
| --- | --- | --- | --- | --- |
| Anti-Nestin Antibody, Clone Rat-401 | Millipore Sigma | MAB353 | Mouse | 1:200 |
| Anti-Tubulin β3 (TUBB3) Antibody | BioLegend | 801213 | Mouse | 5 μg/mL |
| Anti-GFAP Antibody | Sigma-Aldrich | SAB4300647 | Rabbit | 1:500 |
| Anti-O1 Antibody | Millipore Sigma | MAB344 | Mouse | 20 μg/mL |
| Anti-Myelin Basic Protein (MBP) Antibody | Sigma-Aldrich | M3821 | Rabbit | 1:200 |
| Anti-O4 Antibody, Clone 81 | Millipore Sigma | MAB345 | Mouse | 20 μg/mL |
| Anti-CNPase Antibody, clone 11-5B | Millipore Sigma | MAB326 | Mouse | 10 μg/mL |
| Goat Anti-Mouse IgG (H+L) Cross-Adsorbed Secondary Antibody, Alexa Fluor 488 | ThermoFisher | A-11001 | Goat | 1:500 |
| Goat Anti-Rabbit IgG (H+L) Cross-Adsorbed Secondary Antibody, Alexa Fluor 555 | ThermoFisher | A-21428 | Goat | 1:500 |

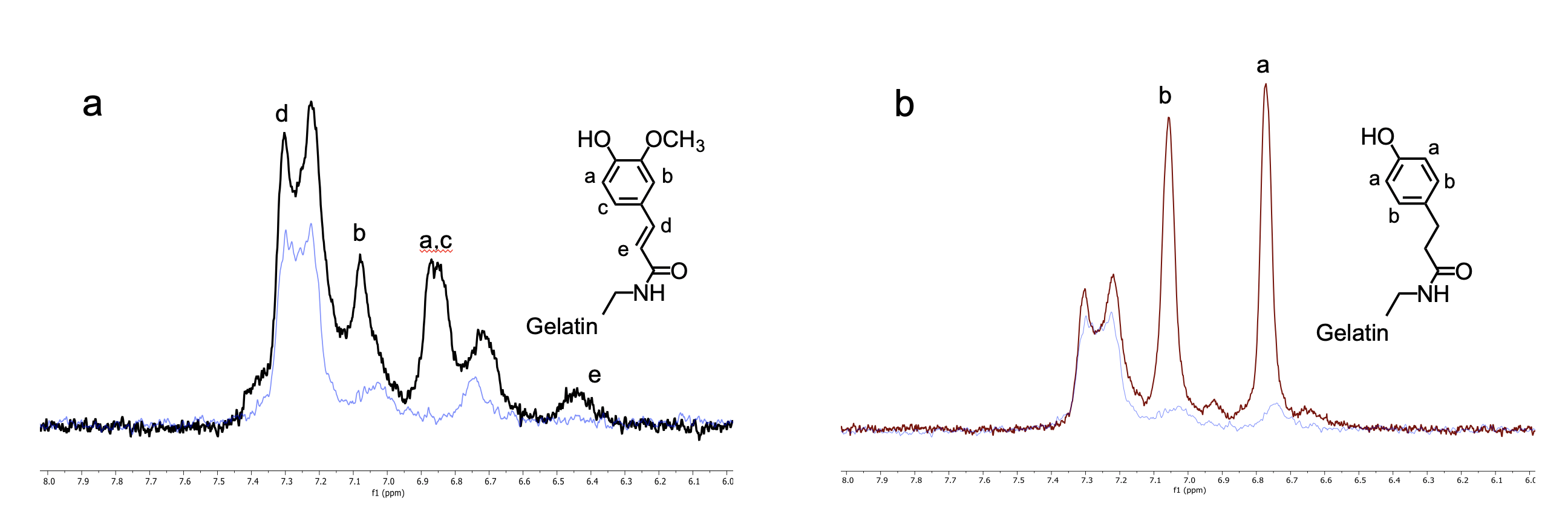

**Figure S1.** NMR characterization of gelatin-ferulic acid (GelFA) and gelatin-hydroxyphenylpropionic acid (GelHPA) conjugates. (a) ^1^H NMR spectra of GelFA (black) and gelatin (blue) (500 MHz, D_2_O, rt): δ 7.35 – 7.28 (m, 1H), 7.12 – 7.00 (m, 1H), 6.95 – 6.79 (m, 2H), 6.51 – 6.40 (m, 1H). (b) ^1^ H NMR spectra of GelHPA (red) and gelatin (blue) (500 MHz, D_2_O, rt): δ 7.11 – 7.00 (m, 2H), 6.82 – 6.70 (m, 2H).

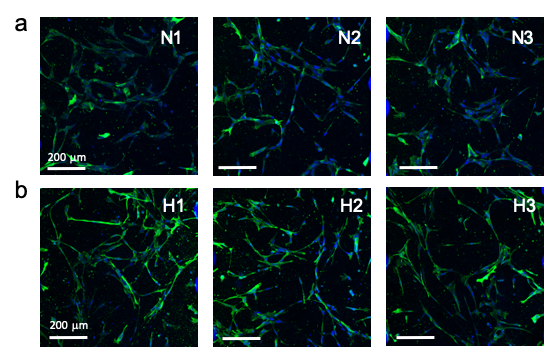

**Figure S2. Glial fibrillary acidic protein (GFAP) expression in astrocytes.** Normal human astrocytes cultured in 2wt% GelHPA hydrogels in (a) normoxia and (b) hypoxia (1% O_2_) for 24h. GFAP (green) and Hoechst (blue, nuclei).
